# CD6 facilitates canonical type I interferon signaling in T cells

**DOI:** 10.64898/2026.11.24.768990

**Authors:** Amber Cardani-Boulton, Adam M. Boulton, Ruiting Zhou, Rachel G. Scheraga, Brian T. Gaudette, Feng Lin, Yuxin Wang, Cornelia C. Bergmann

**Author notes:** Address correspondence to Dr. Bergmann,.

## Abstract

The T cell scavenger receptor CD6 is a widely recognized positive and negative modulator of T cell receptor signaling. Although CD6 is therapeutically targeted the mechanisms underlying its versatile functions and clinical efficacy remain unclear. Herein we investigated CD6 mediated suppression of T cell activation during murine coronavirus infection. Single-cell RNA sequencing revealed impaired upregulation of canonical type I IFN (IFN-I) stimulated genes (ISGs) in CD6 deficient CD4 T cells, which coincided with enhanced T cell activation, proliferation, and T follicular helper differentiation following infection. Direct IFN-I stimulation confirmed that CD6 was required for IFN-I induced STAT1 phosphorylation and supported canonical STAT1/2 dependent ISG upregulation in both murine and human CD4 and CD8 T cells. Additionally, in primary human T cells, CD6 blockade attenuated IFN-I induced cytotoxic effectors and anti-proliferative responses. Mechanistically, cell-surface CD6 functionally associates with the IFNalpha receptors (IFNaR)1 and IFNaR2 to support receptor association, while the cytoplasmic tail of CD6 also interacts with the IFNaR2 cytoplasmic tail and STAT1. These studies identify CD6 as a transmembrane component of the IFNaR signalosome on T cells that facilitates canonical IFN-I signaling, a pathway established to modulate T cell responses across infections, autoimmunity, cancer, and vaccination. Therefore, these findings not only provide additional insight into the mechanisms underlying anti-CD6 therapies but broaden the therapeutic potential of targeting CD6 to manipulate IFN-I controlled T cell activation, proliferation, apoptosis and effector profiles.

**One Sentence Summary:** CD6 is a previously unrecognized component of the IFNaR signaling complex in T cells that facilitates IFNaR1-IFNaR2 organization and canonical IFN-I signaling, revealing a novel immunoregulatory pathway that can be therapeutically targeted to modulate IFN-I dependent T cell activation and cytotoxic effector programs.

## Introduction

CD6 is a therapeutically targeted transmembrane receptor expressed primarily by T cells, but can also be found on some B1a B cells as well as human CD56^+^ NK cells^1–3^. The extracellular domain of CD6 is comprised of three scavenger receptor cysteine rich (SRCR) domains (SRCRd1-3) and a 244 amino acid long cytoplasmic tail that contains numerous docking sites for signaling proteins^4^. As a scavenger receptor, CD6 exhibits some promiscuity, with its most prominently studied extracellular ligands being CD166/ALCAM and CD318/CDCP1, which bind to the CD6 membrane proximal SRCRd3 and membrane distal SRCRd1 domain respectively^4–6^.

Findings that polymorphisms as well as abnormal expression of CD6 and its ligands CD166 and CD318 are associated with susceptibility and severity of multiple autoimmune diseases and cancers has led to clinical use of the monoclonal anti-CD6 blocking antibody Itolizumab^10–16^. In addition to Itolizumab, the anti-CD6 monoclonal antibody UMCD6, which has greater affinity and extended capacity for blocking CD6-ligand interactions, has also shown therapeutic and protective efficacy in experimental tumor and autoimmune models like autoimmune encephalomyelitis and uveitis^13,17–20^.

During T cell activation CD6 incorporates into the T cell receptor (TCR) signalosome, where it acts as a scaffolding protein that can both positively and negatively modulate the TCR signal strength to control T cell activation through incompletely identified mechanisms^1,6–9^. *In vitro* T cells were shown to use CD6 to anchor onto ALCAM expressing antigen presenting cells (APC), thus allowing for prolonged direct contact necessary to initiate T cell proliferation^66^. CD6 further supports T cell activation under anti-CD3/CD28 stimulation and mass spectrometry data of an affinity tagged CD6 pull down demonstrated that CD6 rapidly associated with numerous positive regulators of TCR signaling (Zap70, SLP76, GRB2, GRK6, SHIP1, VAV1, *etc*.) while simultaneously binding to the negative regulator UBASH3a during T cell activation^7^. However, under conditions in which CD6 suppresses T cell activation, CD6 has also been shown to bind to GTPase activating protein of Ras (RasGAP), which in turn limited Ras activation during T cell activation *in vitro*^9,70^. These findings emphasize that CD6 regulation of T cell activation and differentiation is complex, making the identification of mechanisms underlying its plastic functions challenging. Therefore, despite the therapeutic efficacy of anti-CD6 treatments in psoriasis and rheumatoid arthritis patients, the mechanisms determining the context-dependent functions of CD6 remain incompletely defined and cannot be fully explained by the characterized interactions with identified ligands^20^.

As part of our investigation into CD6 we have previously shown that CD6-m CD4 T cell activation and T follicular helper cell (Tfh) differentiation during murine coronavirus (mCoV) encephalomyelitis^113,17^. In this report, investigation into the mechanisms underlying CD6 suppressive functions in the mCoV model revealed that CD6 enables canonical type I interferon (IFN-I) signaling in human and murine T cells. IFN-Is are a class of pleiotropic cytokines that all bind to a heterodimeric receptor composed of the IFNα receptor 1 and 2 (IFNaR1 and IFNaR2). Canonically, IFN-I engagement results in activation of the receptor-associated kinases TYK2 and JAK1, which in turn leads to the phosphorylation of STAT2 and STAT1. Complexed with IRF9, phosphorylated STAT1 and STAT2 (pSTAT1 and pSTAT2) form the ISGF3 which initiates the transcription of hundreds of interferon-stimulated genes (ISGs) by binding to DNA ISREs^31^. While prominently known for their innate antiviral functions, IFN-Is also exhibit versatile control over T cell activation, proliferation, effector differentiation, and apoptosis^21–23^. Here, we show that extracellular CD6 functionally associates with the IFNaR signaling complex and facilitate organization of the receptor chains, while intracellularly CD6 associates with IFNaR2 and STAT1. These findings have broad implications as they indicate that CD6 may be targeted to either mitigate or enhance T cell ISGs transcriptional programs in various clinical settings^21–23^.

## Results

### CD6 deficient CD4 T cells have impaired ISG upregulation during mCoV infection

To probe how CD6 regulates CD4 T cell activation and differentiation during mCoV encephalomyelitis^1^, we performed single-cell sequencing (scSeq) of negatively enriched CD4 T cells isolated from the CNS draining cervical lymph nodes (dLNs) of WT and CD6 KO mice on day 4 post infection (PI) (Supplemental Figure 1A). Cell populations were plotted by a UMAP and manually identified using established markers (Figure 1A-B) revealing clusters of T regulatory (Treg) cells (*Foxp3^+^*), Tfh cells (*Cxcr5^+^* and *Il21^+^)*, T helper 1 (Th1) cells (*Tbx21^+^* and *Ifng^+^*), proliferating cells (*Mki67^+^*), and a cluster that was heterogenous for multiple activation markers with an expression profile indicative of active TCR signaling (*Nr4a1^+^*, *Il2ra^+^*, and high *Irf4*)^1^ (Figure 1B and Supplemental Figure 1B). As expected, the vast majority of T cells at day 4 PI had a naïve signature. Consistent with our published flow cytometry data CD6 KO CD4 T cells had increased activation (*Cd44^+^*), Tfh differentiation, and proliferation, but no difference in Treg or Th1 differentiation (Figure 1C)^1^.

**Figure 1:**
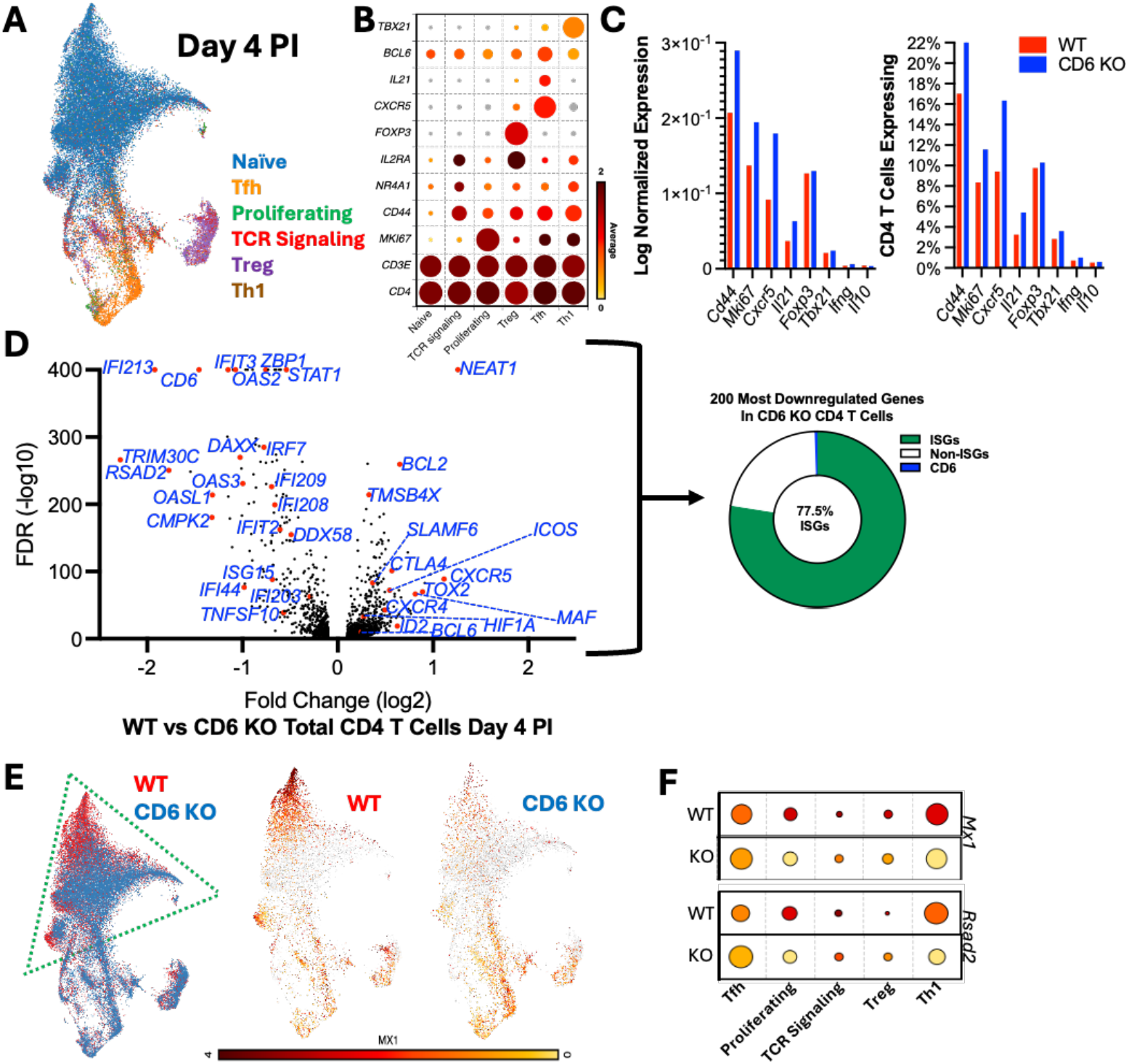
CD6 deficient CD4 T cells have impaired ISG upregulation during mCoV infection. scSeq analysis of CD4 T cells isolated from the dLN of mCoV infected WT and CD6 KO mice on 4 days PI showing **A)** UMAP clustering, **B)** bubble heatmap of genes identifying distinct T helper subsets, **C)** expression of select CD4 T effector genes, and **D**) a volcano plot of WT versus CD6 KO DEGs in CD4 T cells. **E)** UMAP comparing WT and CD6 KO CD4 T cells on day 4 PI (left) and highlighting *Mx1* expression (right). Green triangle highlighting the naïve cells. **F)** Bubble heatmap showing *Mx1* and *Rsad2* expression on day 4 PI separated by CD4 T cell cluster.

Differentially expressed genes (DEG) between all WT and CD6 KO CD4 T cells at day 4 PI revealed a dramatic deficit in expression of STAT1/STAT2-mediated ISGs in the CD6 KO CD4 T cells (Figure 1D). In fact, >77% of the top 200 significantly downregulated genes in CD6 KO CD4 T cells were ISGs. Conversely, several STAT3 target genes (*Neat1*, *Bcl2*, *Socs3*, and *Bcl6*) were significantly upregulated in CD6 KO CD4 T cells^24–28^ (Figure 1D), in line with enhanced Tfh differentiation in mCoV infected CD6 KO mice^1^. Analysis of which CD4 T cell populations expressed ISGs at day 4 PI revealed that the naïve and early TCR signaling clusters exhibited the largest difference in ISG expression between the WT and CD6 KO CD4 T cells (Figure 1E outlined by green dashed line), but that all CD6 KO CD4 T cell subsets had diminished ISG expression (Figure 1F). As T cells have diminished capacity for STAT1-mediated IFN-I signaling after TCR activation^62^, separating CD4 T cells by activation state (defined here by CD44 expression) revealed that WT CD44^-^CD4 T cells displayed the highest ISG expression levels while the CD44^-^CD6 KO T cells had the lowest (Supplemental Figure 1C).

### CD6 deficient T cells exhibit largely preserved homeostatic features

To control for any preexisting homeostatic differences in CD6 KO T cells, we simultaneously performed scSeq on CD4 T cells from uninfected (UI) WT and CD6 KO mice, which showed no obvious differences including in CD4 T cell clustering (Supplemental Figure 2A-C). Notably the scSeq data showed similar transcript levels for the IFNaR signaling components and *Ifit1*, a strongly IFN-I induced ISG, between WT and CD6 KO CD4 T cells from UI mice (Supplemental Figure 2D). Flow cytometry analysis confirmed comparable IFNaR1 protein levels on the surface of CD4 and CD8 T cells from UI WT and CD6 KO mice (Supplemental Figure 2E). These data supported that reduced ISG upregulation in CD6 KO CD4 T cells was a direct result of signals induced by viral infection.

### CD6 supports IFN-I induced ISG upregulation in murine T cells

To determine if CD6 facilitates IFN-I signaling independent of infection, splenic CD4 T cells from UI WT and CD6 KO mice were negatively enriched and stimulated *ex vivo* with IFNβ. Analysis of ISG expression by qPCR after 4 hours of stimulation confirmed IFN-I induced ISG upregulation was impaired in CD6 KO CD4 T cells (Figure 2A). IFNβ was also administered by IP injection to WT and CD6 KO mice. FACS isolated splenic CD4 and CD8 T cells from CD6 KO mice again showed impaired ISG upregulation by qPCR compared to WT mice (Figure 2B).

**Figure 2:**
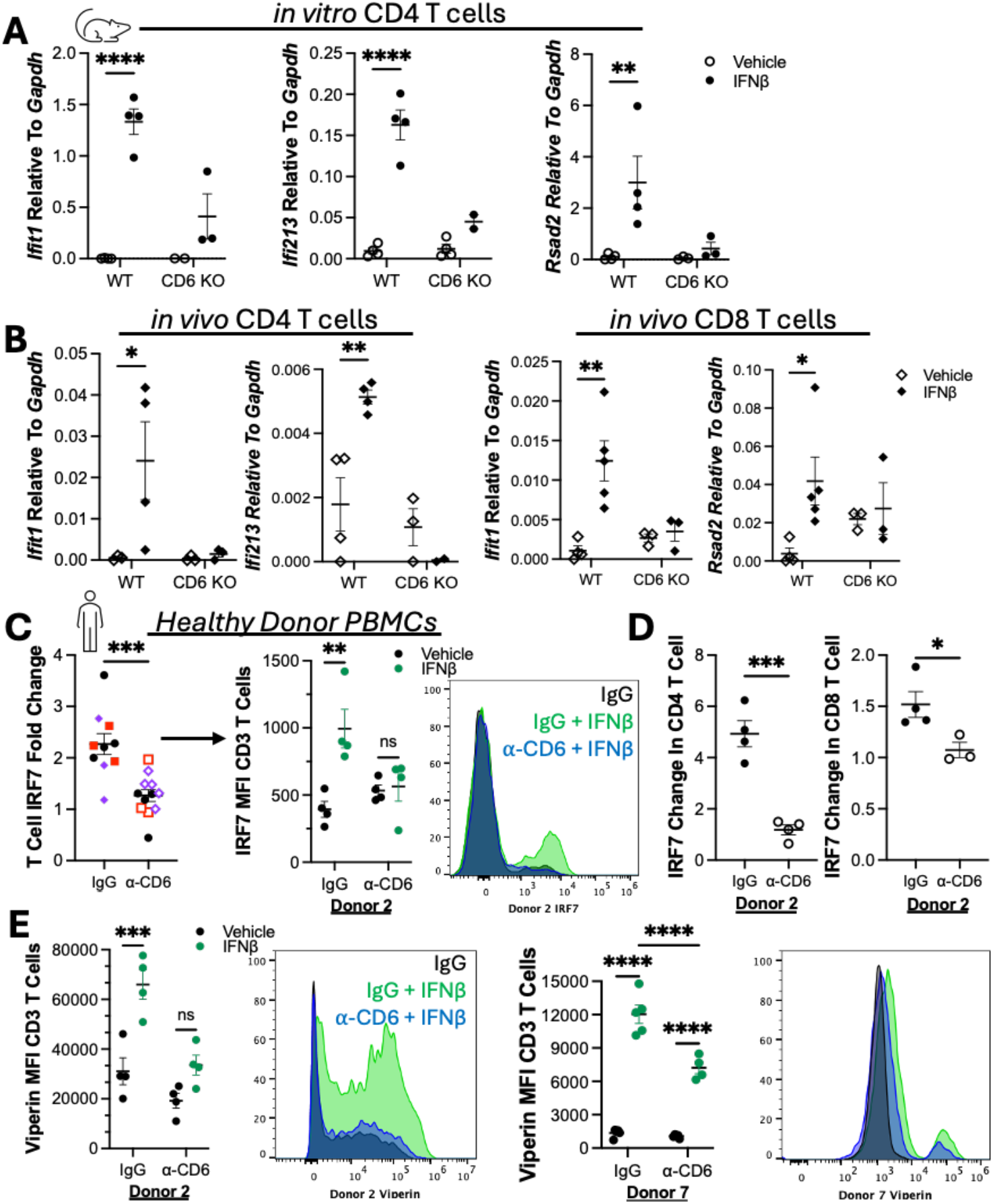
CD6 is required for optimal upregulation of ISG programs in T cells. ISGs measured by qPCR in **A**) splenic CD4 T cells after 4 hours of *ex vivo* IFNβ stimulation and in **B**) splenic CD4 (left) and CD8 (right) T cells ∼18hours after *in vivo* IFNβ stimulation. Circles represent an individual mouse whose T cells were used across experimental conditions. Each diamond represents an individual mouse from two pooled experiments. **C)** Human PBMCs treated with IgG control or anti-CD6 followed by co-stimulation with IFNβ overnight were analyzed for upregulation of the ISGs IRF7 by flow cytometry. Individual donors are denoted by color and data were combined by normalizing as fold change (IFNβ stimulation/ Vehicle control). Median IFNβ responder Donor 2 (black circles) is shown as representative for IRF7. **D**) CD3 T cells for Donor 2 separated into CD4 and CD8 T cells. **E**) The ISG Viperin was similarly analyzed by flow cytometry. All data is graphed using standard error mean. Data were analyzed in **C** and **E)** using 2-way ANOVA with a Bonferroni post-test, or for fold-change **C** and **D**) in which was analyzed using a two-tailed unpaired student’s T-test.

### CD6 blocking antibody UMCD6 can limit ISG upregulation in human T cells

We next tested if CD6 blockade could be used to modulate ISG upregulation in human T cells by stimulating PBMCs from healthy donors with IFNβ while blocking CD6^13,17–19^. Given the unknown nature underlying CD6-mediated IFN-I signaling, the CD6 blocking antibody UMCD6 was selected for our analysis due to its higher binding affinity, slower dissociation kinetics, and greater ability to interfere with CD6-CD166 interactions compared with Itolizumab^20^. Flow cytometry analysis confirmed that control treated human T cells upregulated the ISG IRF7 in response to IFNβ, but that CD6 blockade significantly impaired upregulation across multiple donors (Figure 2C). Notably, UMCD6 treatment impeded the IRF7 upregulation in both CD4 and CD8 T cells from healthy donors (Figure 2D). Upregulation of the ISG Viperin was also significantly impaired (Figure 2E). These data demonstrate that CD6 plays a crucial role in IFNβ induced ISG upregulation in both human and murine CD4 and CD8 T cells.

### CD6 functionally associates with the IFNaR signaling complex, promotes receptor organization, and supports canonical IFN-I responses

As a CD6 blocking antibody was able to impair IFN-I mediated ISG upregulation, we next asked if supplementing CD6 could increase ISG responses. We therefore added the extracellular domain of CD6, also referred to as soluble CD6 (sCD6)^41–42^, to the CD6 negative murine fibroblast LL171 cell line, which stably expresses an IFN-I Stimulated Response Element (ISRE)-luciferase construct as a reporter of canonical IFN-I signaling^43^. Overnight IFNβ stimulation strongly induced luciferase production in LL171 cells, which was significantly enhanced in the presence of sCD6 (Figure 3A). Importantly, sCD6 treatment alone did not induce luciferase production (Figure 3A), demonstrating that extracellular CD6 can amplify IFN-I induced ISG upregulation.

**Figure 3:**
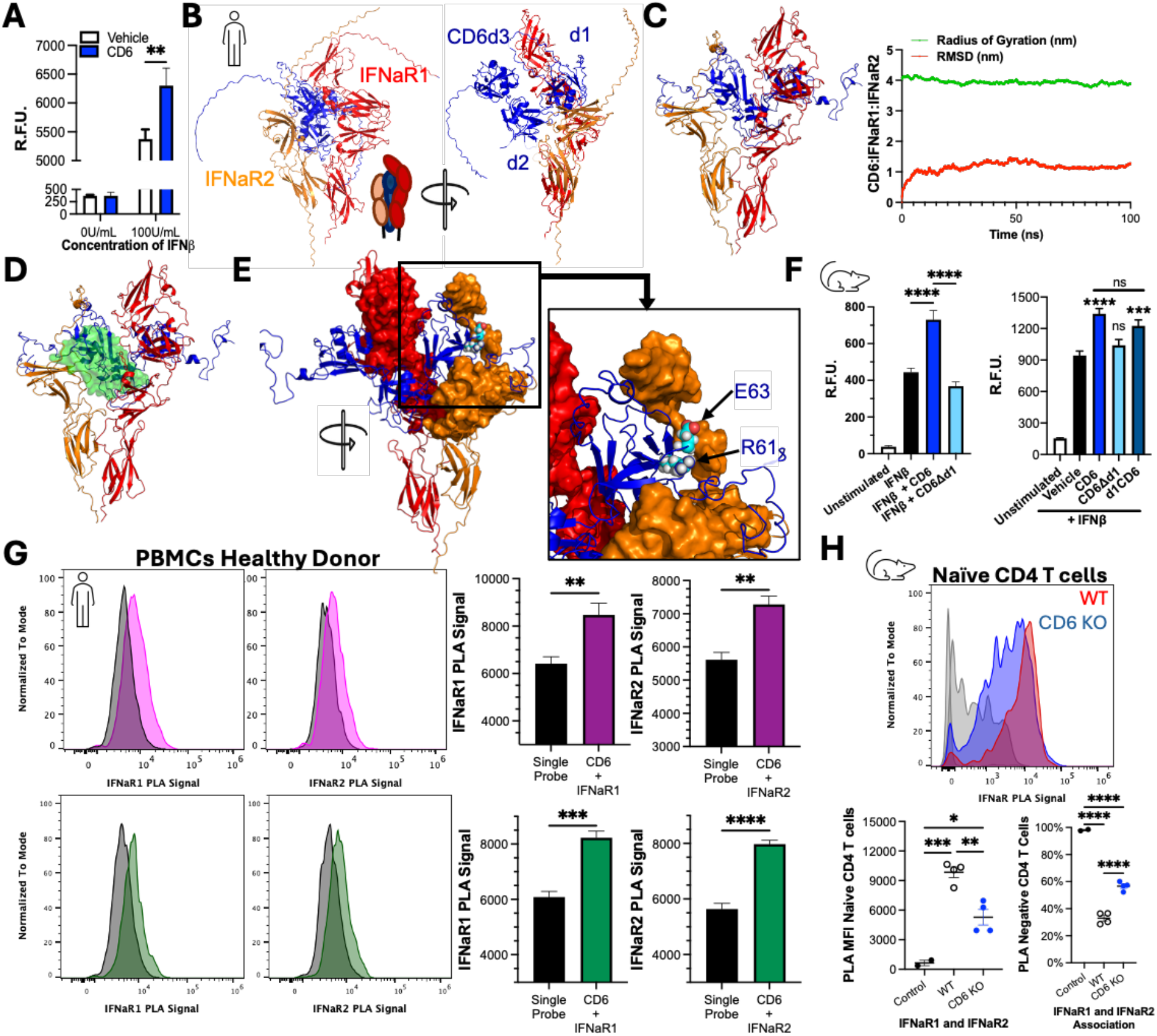
CD6 associates with the IFNaR complex on the cell surface to amplify IFN-I induced transcription. **A)** LL171 cells treated with CD6 and then co-stimulated with IFNβ. After overnight incubation luciferase was measured as relative fluorescence units (R.F.U.). **B)** AlphaFold predicted structure of CD6 (blue) with IFNaR1 (red) and IFNaR2 (orange). **C)** GROMACS analysis of interactions between CD6 (blue), IFNaR1 (red) and IFNaR2 (orange) including molecular dynamics over 100ns simulation. **D)** Alignment of the solved IFNaRs with IFNalpha2 (green) bound (PDB #SE3) over the energy minimized model of the CD6-IFNaRs prediction. IFNalpha2 is represented using surface modeling to show it occupies the same space as the CD6 SRCRd1. **E)** Surface fill of IFNaR1 and IFNaR2 residues predicted to be within 20Å of the CD6 SRCRd1. CD6 R61 and E63 sidechains are modeled using spherical representation and colored by element (red= nitrogen, white= hydrogen, and cyan= carbon). **F)** LL171 cells treated with CD6Δd1 or d1 of CD6 at 100nM and then co-stimulated with IFNβ. After overnight incubation luciferase was measured as relative fluorescence units (R.F.U.). **G)** Flow cytometry PLA fluorescent signal generated by oligonucleotide conjugated antibodies recognizing anti-IFNaR1 or anti-IFNaR2 along with anti-CD6 on unpermeablized human PBMCs. **H)** Naive splenic CD4 T cells from WT and CD6 KO mice were analyzed for IFNaR1 and IFNaR2 proximity by PLA fluorescence. Quantification of PLA MFI (right) and percent of CD4 T cells negative for PLA signal (left). Each data point represents CD4 T cells from an individual mouse. Data were graphed and analyzed in GraphPad PRISM using a 2-way (**A**) or 1-way ANOVA (**F and H**) with a Bonferroni post-test or an unpaired students T test (**G**).

Having established that sCD6 was sufficient to enhance IFN-I induced ISG upregulation we next assessed a potential interaction between CD6 and cell surface IFNaRs by querying AlphaFold^35–37^, which predicted that the CD6 SRCRd1 interacts with IFNaR1 and IFNaR2 (Figure 3B). Analysis of predicted CD6-IFNaR ternary complex over a 100-nanosecond simulation using the molecular dynamics software GROMACS^38–40^ corroborated that the interaction was stable as measured by the consistency of the root mean square deviation (RMSD/ red line) and radius of gyration over time (Figure 3C). Importantly, the receptor ternary complex did not dissociate throughout the simulation. Intriguingly, overlay of the solved IFNalpha2-IFNaR structure (PDB-3SE3) showed the SRCRd1 of CD6 was predicted to interact with the IFNaRs by occupying the IFN-I binding pocket^63^ (Figure 3D). Further examination revealed numerous IFNaR1 and IFNaR2 residues within 20Å of the CD6 SRCRd1, the domain that both Itolizumab and UMCD6 bind, indicating there would be steric clash with the blocking antibodies (Figure 3E). Indeed, examination of CD6 R61 and E63 which are required for Itolizumab binding and located in a region that overlaps the UMCD6 binding epitope, reveals numerous IFNaR2 residues located within 20Å of these two residues (Figure 3E).

As the SRCRd1 of CD6 was predicted to be the predominant interaction site with the IFNaR chains, we next tested the necessity of SRCRd1 to support IFN-I signaling in LL171 cells. Unlike full length sCD6, sCD6 lacking the SRCRd1 (CD6Δd1) did not increase IFNβ induced luciferase production (Figure 3F). Conversely, the SRCRd1 alone was sufficient to support canonical IFN-I signaling, but with less consistency than full length sCD6 (Figure 3F). These data demonstrate that extracellular CD6 supports IFN-I responses through its SRCRd1, likely through direct interactions.

We therefore set out to confirm that CD6 associates with the IFNaRs on the surface of untransformed primary human T cells using a DuoLink flow cytometry-based Proximity Ligation Assay (PLA), which generates fluorescence when antibodies recognizing two distinct proteins are within ∼40nm of each other^44–45^. Antibodies outside of the predicted SRCRd1-IFNaR1/2 interaction interface were utilized with the understanding that the distance between binding epitopes would be greater (CD6 SRCRd3 binding OX124 coupled with either anti-IFNaR1 clone 230549B1 or anti-IFNaR2 clone EPR24706-26)^20^. Indeed, CD6 association with the IFNaR1 as well as IFNaR2 could be detected on primary non-permeabilized human PBMCs (Figure 3G), despite the low levels of IFNaR1 and IFNaR2 expression on human T cells^61^ (Supplemental Figure 2E).

Given that CD6 occupies the IFN-I binding pocket, we next tested if CD6 promotes organization of the IFNaR1-IFNaR2 heterodimeric receptor complex by measuring surface IFNaR1 association with IFNaR2 on naïve murine WT and CD6 KO splenic CD4 T cells by PLA. Indeed, IFNaR1-IFNaR2 association was significantly diminished on CD6 KO naïve CD4 T cells (Figure 3H), with IFNaR1-IFNaR2 association falling below detection levels on an average of 56.6% of CD6 KO CD4 T cells (Figure 3H). Collectively, these data support that CD6 supports the organization and association of IFNaR1 and IFNaR2 on T cells and facilitates canonical IFN-I signaling.

### The cytoplasmic tail of CD6 interacts with the intracellular domain of IFNaR2 and STAT1

Given that IFNaR dimerization and canonical IFN-I signaling occurs in numerous cell types that are CD6 negative and that some CD6 KO T cells still have IFNaR1-IFNaR2 association comparable to WT T cells, CD6-mediated IFNaR organization is unlikely to fully explain CD6-mediated canonical IFN-I signaling in T cells. Based on the findings that CD6 is a scaffolding protein for the TCR^8, 64^, JAK1 co-immunoprecipitated with CD6^30^, and CD6 promotes STAT1 phosphorylation during TCR activation in human T cells^30, 64^, we probed whether CD6 might similarly support IFN-I signaling as a component of the intracellular IFNaR signalosome.

First, to test if CD6 supports signaling through the IFNaR complex, the ability of CD6 KO CD4 T cells to phosphorylate STAT1 (pSTAT1) in response to IFN-I was examined. Splenic CD4 T cells negatively enriched from UI WT and CD6 KO mice were stimulated with IFNβ *ex vivo* for 15 minutes and pSTAT1 was measured by flow cytometry. Unlike WT CD4 T cells, CD6 KO CD4 T cells did not phosphorylate STAT1 in response to IFNβ, even at high doses, despite showing elevated baseline pSTAT1 levels (Figure 4A). These results linked impaired ISG upregulation in CD6 KO T cells to an upstream defect in IFN-I signaling at the receptor.

**Figure 4:**
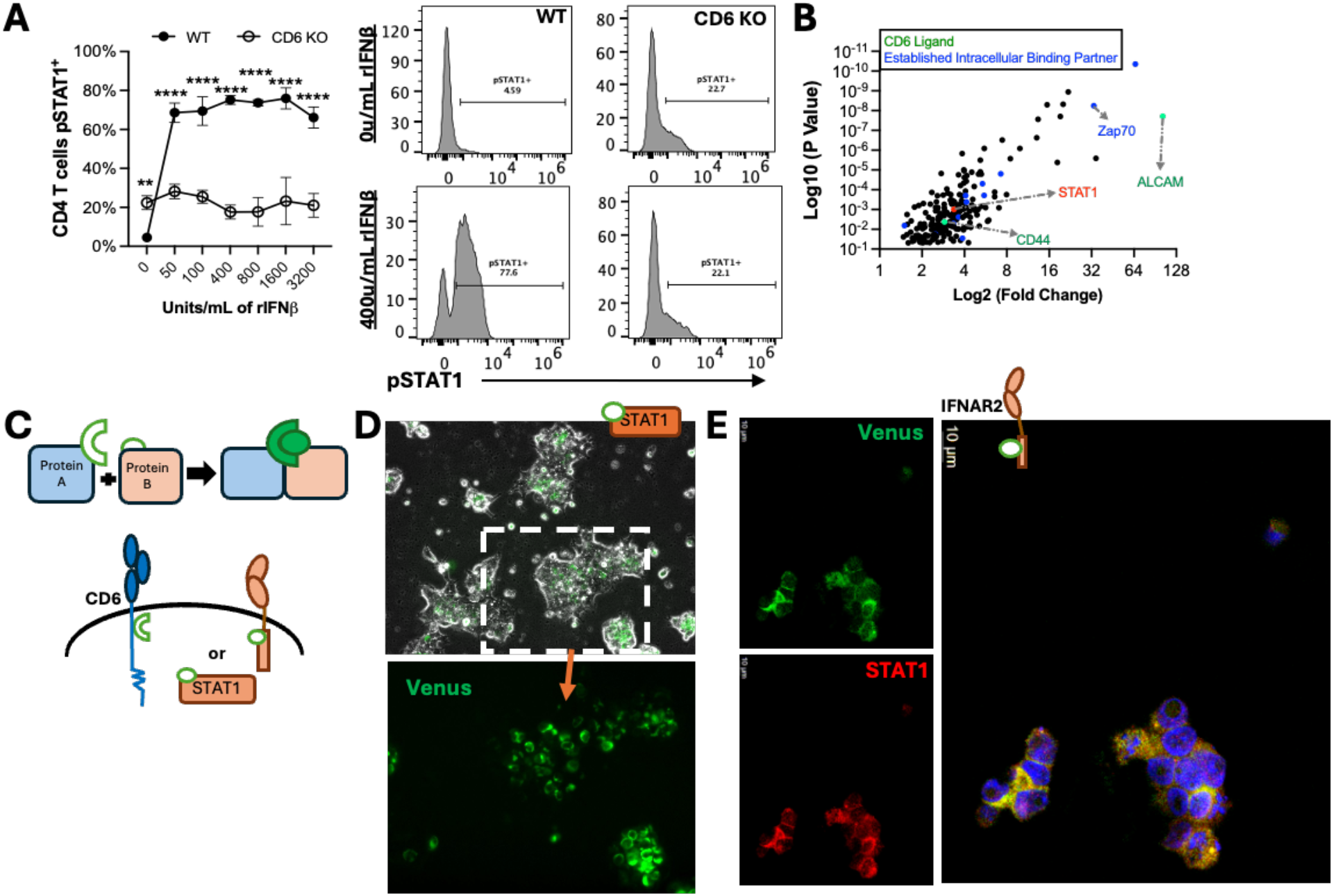
The cytoplasmic tail of CD6 interacts with the intracellular domain of IFNaR2 and STAT1. **A)** pSTAT1 was monitored in splenic CD4 T cells treated with IFNβ for 15min *ex vivo* by flow cytometry; representative histograms are shown on right. **B)** Proteins from a MassSpec analysis from a pull-down of CD6 performed on murine CD4 T cells as done by Mori *et al.*^2^ Established CD6 ligands (green) and CD6 binding partners (blue) as well as STAT1 (red). **C)** BiFC experimental design for **D)** CD6 Venus-N and STAT1 Venus-C transfected HECK293T cells with brightfield (top) and Venus/GFP signal as well as **E)** HECK293T cells stably expressing CD6 Venus-N and IFNaR2 Venus-C and stained with anti-STAT1. Images are representative of at least 3 replicates. Data were graphed in GraphPad Prism using standard error mean and analyzed using a 2-way ANOVA with a Bonferroni post-test (**A**) or as described in method section (**B**).

Next, Mori *et al.’*s proteomics data set of a CD6 co-immunoprecipitation from murine CD4 T cells that had been transduced *ex vivo* with affinity-tagged CD6 was analyzed^7^. While Mori *et al.* focused on CD6 binding partners that increased during TCR stimulation to probe how CD6 modulated TCR signaling, our analysis focused on constitutive CD6 binding partners (without TCR stimulation). As expected, T cell expressed extracellular CD6 ligands (ALCAM and CD44) as well as some of its established intracellular binding partners pulled down with CD6 (Figure 4B). Additionally, we found STAT1 significantly bound to CD6, confirming STAT1 as a CD6 binding partner (Figure 4B). The IFNaRs were not detected in this dataset, which was not surprising as these T cells had undergone TCR-mediated activation and long-term culture during transduction and TCR activation is known to diminish T cell capacity for IFN-I singling^62^. Additionally, neither the tyrosine receptor kinases TYK2 and JAK1, nor STAT2 were identified as significantly enriched in the CD6 pull-down^7^ despite JAK1 having been shown to co-immunoprecipitate with CD6^30^. Therefore, STAT1 binding to CD6 was confirmed using Bimolecular Fluorescence Complementation (BiFC) assays in which two non-fluorescent fragments of the fluorescent protein Venus are fused to two separate proteins of interest. Direct interactions between these tagged proteins results in the Venus fragments fusing, thus generating Venus/GFP fluorescence^32–33^ (Figure 4C). HEK293T cells expressing human CD6Venus-N and STAT1Venus-C revealed a strong Venus signal (Figure 4D), supporting that the cytoplasmic tail of CD6 interacts with STAT1, which is also consistent with published studies showing decreased pSTAT1 levels during TCR stimulation when CD6 is inhibited^30^.

As STAT1 associates with multiple receptors, we further used BiFC to establish that CD6 operates intracellularly at the IFNaR signaling complex by probing for CD6 interactions with the cytoplasmic tail the IFNaR2 chain, the primary platform of IFN-I signaling^31,34^. HEK293T cells stably expressing human CD6Venus-N and human IFNaR2Venus-C demonstrated a direct interaction between the cytoplasmic tails of CD6 and IFNaR2 (Figure 4E). Co-staining with anti-STAT1 confirmed that endogenous STAT1 localized to the CD6-IFNaR2 complex (Figure 4E). Collectively, these data support a model in which CD6 is a component of the IFNAR signaling complex in T cells. Its extracellular association with IFNaR1 and IFNaR2, together with intracellular interactions with IFNaR2 and STAT1, provides a mechanistic framework for how CD6 facilitates signaling through the IFNaR complex to promote STAT1 phosphorylation and ISG upregulation.

### CD6-mediated regulation of human T cell activation and proliferation differs under IFN-I conditions

To determine how IFN-I signaling impacts effector functions we analyzed a publicly available bulk RNAseq time course of human CD4 and CD8 T cells activated *in vitro* in the presence or absence of exogenous IFNβ (GSE195543)^46^. DEG analysis demonstrated a strong ISG response as early as 4 hours post IFNβ treatment in both CD4 (Figure 5A) and CD8 (Figure 5B) T cells. Additionally, *TNFSF10* encoding the death receptor TRAIL^47^ was highly upregulated at 4 hours in both CD4 and CD8 T cells, while *MKI67* expression was significantly decreased in IFNβ treated CD8 T cells (Figure 5A-B; left panel). After 72 hours of IFNβ stimulation CD8 T cells, and to a lesser extent CD4 T cells, exhibited significantly elevated expression of the cytotoxic effectors *FASLG*, *GZMB*, *PRF1*, and *TNSF10* (Figure 5A-B; left column), demonstrating that IFN-I signaling induces cytotoxic effectors in human CD4 and CD8 T cells.

**Figure 5:**
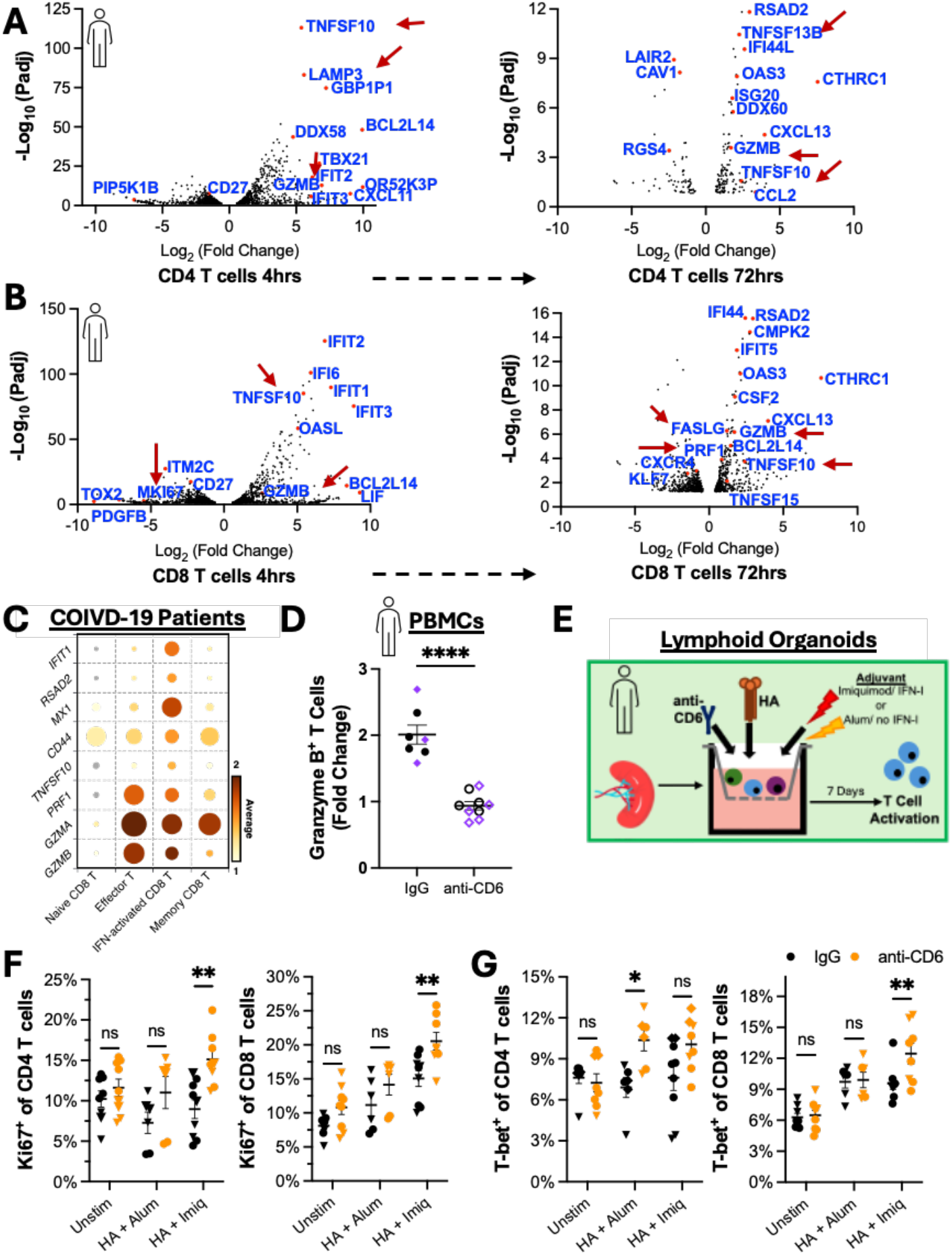
CD6-mediated regulation of human T cell activation and proliferation differs under IFN-I conditions. Volcano plot of DEGs from a bulk RNA-seq data set of human **A)** CD4 and **B)** CD8 T cells activated *in vitro* with and without exogenous IFNβ after 4 and 72 hours (GSE195543). **C)** Bubble heatmap analyzing cytotoxic effectors in T cells from patients with COIVD-19 in GSEA 155223. **D)** Granzyme B upregulation in T cells from human PBMCs following overnight stimulation with IFNβ measured by flow cytometry. **E)** Schematic of lymphoid organoid experimental workflow. **F)** Flow cytometry analysis of T cell proliferation and **G**) T-bet expression in human organoids after 7 days in culture with indicated treatments. Organoids were generated from 2 donors (circles vs triangles). Each donor was used in duplicate experiments to ensure reproducibility. Data are graphed at standard error mean and analyzed using a **D)** 2-tailed unpaired T-test or **F-G)** 2-way ANOVA with Bonfernoi post-test.

The *in vivo* significance of canonical IFN-I signaling in T cells during viral infections was examined by analyzing an ISG high CD8 T cell subset identified in Unterman *et al.*’s scSeq data from COVID-19 patients (GSEA 155223)^48–49^. This cluster also showed high levels of Granzyme A and B compared to naïve and memory CD8 T cells as well as significantly higher TRAIL expression, consistent with the ability of IFN-Is to directly upregulate TRAIL (Figure 5C) ^47^.

We therefore tested the ability of CD6 to support IFN-I induced cytotoxic effectors in human T cells. To first separate the role of CD6 in regulating IFN-I signaling from its role in modulating TCR signaling, PBMCs from healthy donors were stimulated with IFNβ alone in the presence of CD6 blocking antibody UMCD6. Even without TCR activation IFNβ modestly increased Granzyme B expression, which was prevented by CD6 blockade (Figure 5D). Therefore IFN-I driven expression of cytotoxic effectors in human T cells can be dampened by anti-CD6 blocking antibodies.

Lastly, to determine whether the enhanced proliferation and T cell accumulation observed in the mCoV infected CD6 KO mice^1^ (Figure 1C) could be recapitulated in a human system, we examined human T cell activation in splenic-derived human lymphoid organoids^65^. Lymphoid organoids were stimulated with recombinant influenza A virus HA (A/CA/07/2009) plus either Imiquimod (Imiq) or Alum adjuvant to compare IFN-I inducing to non-IFN-I inducing conditions^65, 67–69^ (Figure 5E). Similar to mCoV infected CD6 KO mice, UMCD6 treated human CD4 T cells underwent increased proliferation in IFN-I inducing condition but not with Alum adjuvant (Figure 1C and 5F). This finding extended to UMCD6 treated human CD8 T cells, suggesting an anti-proliferative role of CD6 specifically in IFN-I inducing viral settings (Figure 5F). T-bet expression in UMCD6 treated human CD4 T cells was not altered under IFN-I inducing conditions, again recapitulating findings in the mCoV infection model (Figure 1C and 5G). Intriguingly though, CD4 T cell T-bet expression was increased by UMCD6 treatment in the non-IFN-I inducing Alum condition. Conversely, UMCD6 treatment only increased CD8 T cell T-bet expression in IFN-I inducing conditions (Figure 5G). Taken together, these data show CD6 blockade in human lymphoid organoids mimics the enhanced CD6 KO CD4 T cell proliferation found during mCoV infection, but only under IFN-I inducing conditions. Additionally, while we only saw transient CD8 T cell accumulation in lymphoid tissue during mCoV infection, UMCD6 treatment also enhanced human CD8 T cell proliferation and T-bet expression under IFN-I inducing conditions, but did not have a significant impact in non-IFN-I conditions. This data therefore suggests that the presence of IFN-Is is one of the determinants of the plastic outcomes of CD6-mediated regulation of T cell activation in human T cells.

## Discussion

Despite the clinical importance of CD6, molecular insights into its functions are limited and have largely centered on its role in modulating TCR signaling during T cell activation^1–7^. Here, our findings demonstrate that CD6 enables canonical IFN-I signaling in both human and murine CD4 and CD8 T cells, even in the absence of TCR engagement. Direct association of CD6 with the IFNaR on the cell surface was demonstrated by PLA as well as enhanced canonical IFN-I signaling with exogenous sCD6. Significantly, IFNaR1 and IFNaR2 association was undetectable by PLA on more than half of naïve CD6 KO CD4 T calls. Additionally, we identify intracellular interactions between the cytoplasmic tail of CD6 and IFNaR2 as well as STAT1. These data thus establish multiple points of interaction between CD6 and the IFNaR signaling complex and suggest a model whereby CD6 promotes organization of the IFNaR signalosome in T cells to facilitate canonical STAT1-mediated IFN-I signaling. The ability of CD6 to promote IFNβ induced STAT1 phosphorylation and ISG expression in naïve T cells establishes an unrecognized immunoregulatory mechanism contributing to T cell activation, differentiation, effector functions, and pathogenicity and thus has broad basic biological and clinical significance.

Importantly, STAT1-mediated IFN-I signaling is known to shape antiviral T cell responses as well as to promote anti-proliferative, pro-apoptotic, and cytolytic differentiation programs upon subsequent TCR activation^21–22^. Bulk RNAseq analysis confirmed that IFN-I stimulation promoted cytotoxic effector profiles in human T cells (GSE195543)^46^, which we could limit by treating with an anti-CD6 blocking antibody. In the context of mCoV infection, we observed increased T cell proliferation, activation, and accumulation in infected tissue in the absence of CD6^1^. Additionally, increased CD4 Tfh differentiation in dLN of infected CD6 KO mice was accompanied by increased STAT3 regulated transcripts. This observation aligns with previous reports that IFN-I stimulation of STAT1 KO T cells drives STAT3 activation^21^ and that the balance between STAT1 and STAT3 activation during the first 48-hours of LCMV infection governs CD4 T cell commitment between Th1 and Tfh effector fates respectively^50^. Intriguingly, blockade of CD6 in human lymphoid organoids also lead to increased T cell proliferation and CD8 T cell T bet expression under IFN-I inducing conditions. Together, these data emphasize the potential biological significance of CD6 in regulating the balance of T cell effector fates.

The capacity of IFN-I to shape T cell fate across diverse immunological settings, including tumor immunity^51–53^, chronic infection^54–56^, autoimmunity^57^, and vaccination^23^ underscores the potential translational relevance of CD6-facilitated IFN-I signaling. For example, recent work by Wang *et al*. demonstrated that patients with systemic lupus erythematosus and antibody-associated vasculitis have an expanded population of circulating T cells characterized by elevated ISG expression and enhanced cytotoxic effector programs^57^. As both altered CD6 expression and dysregulated IFN-I signatures have been linked to clinical autoimmune diseases such as multiple sclerosis, systemic lupus erythematosus, and lupus nephritis^58–59^, these findings raise the possibility that altered CD6-dependent IFN-I signaling could potentially contribute to clinically pathogenic T cell states. These observations further suggest that therapeutic CD6 blockade may alleviate clinical pathology and disease, not only by altering TCR signaling, but also by influencing IFN-I driven effector profiles. While not addressed in this study, it is worth noting that it is possible that CD6 blockade may allow for greater T cell activation in chronic conditions where IFN-Is become anti-inflammatory and suppress T cell activation. This would line up with other data showing that CD6 blockade promotes regression of CD318 expressing tumors by promoting cytolytic function in NK and CD8 T cells; IFN-Is were however not assessed as part of that study^3,6,19^.

Our data additionally demonstrates that sCD6 directly enhances IFN-I signaling in cells that do not express CD6, pointing to a potential innate-like function of T cell expressed CD6, whereby the proteolytic cleavage of CD6 from the surface of T cells could potentiate IFN-I responses in non-lymphocytic cells. Intriguingly, our published data demonstrated preferential loss of CD6 from the surface of CD4 T cells in the infected tissue but not the dLN^1^. This data suggests that CD6 shedding may be actively regulated, thus providing temporal and anatomical control over the availability of sCD6 to modulate IFN-I signaling in trans.

It is noteworthy that our mechanistic studies focused on naïve T cells stimulated solely with IFN-I to distinguish CD6 regulation of IFN-I signaling from its role during TCR signaling. However, because STAT1 can also be activated downstream of TCR signaling, dissecting the consequences of CD6-mediated STAT1 activation during IFN-I stimulation versus during TCR stimulation will require further investigation^43, 2–5^. Defining how CD6 coordinates its regulation of both TCR and IFNaR signaling will be important to understand how these pathways integrate to control T cells^60^.

Collectively, our study identifies CD6 as a previously unrecognized regulator of IFN-I signal transduction in T cells through its association with components of the IFNaR signaling complex. Thus, this data identifies an unrecognized mechanism that allows T cell activation, proliferation, and effector differentiation to be regulated in a context dependent manner. This mechanism expands the conceptual framework of CD6 biology beyond the immunological synapse and provides novel insight into how CD6-targeting therapies may exert immunomodulatory effects in autoimmunity, infection, and cancer.

## Materials and Methods

### Mice and Infections

All procedures involving mice were approved by the Institutional Animal Care and Use Committee of Cleveland Clinic and carried out in accordance with the US Department of Health and Human Services Guide for the Care and Use of Laboratory Animals and institutional guidelines. CD6 KO mice on the DBA-1 background were generated by the laboratory of Dr. Feng Lin and maintained under pathogen-free conditions in the Cleveland Clinic Research animal facility^13^. WT controls were also maintained onsite and housed in the same room with occasional supplementation as well as rejuvenation of the breeders with mice purchased from Jackson Laboratory (strain number 000670). As previously reported 6-7 week old gender matched male and female WT and CD6 KO mice were intracranially (IC) infected with 10,000 PFU^1^ of the recombinant mCoV strain MHV-A59 whose non-essential open reading frame of gene 4 had been replaced with enhanced green fluorescent protein and was generously provided by Dr. Das Sarma^1^.

### Human tissue

Human PBMCs from healthy donors were purchased frozen and deidentified from StemCell Technologies (#70025.1) specifically requesting unique donors. Splenic cell suspensions were obtained from postmortem specimens that had been collected for research at Cleveland Clinic. The study was determined to be exempt from Institutional Review Boards (IRB) review because it involved the use of postmortem tissue and did not constitute human subject research under institutional and federal guidelines.

### Isolation of murine CD4 T cells

Single-cell suspensions of spleens from UI WT and CD6 KO mice or dLNs on day 4 PI were obtained by manual disruption using 70um mesh screens (Fisher #22-363-548)^1^. CD4 T cells were negatively enriched using the StemCell Technologies mouse CD4 T cell isolation kit and the EasyEight EasySep Magnet following the manufactures protocol (catalog #19852 and #18103).

### Single Cell Sequencing

Isolated cells were counted and fixed with the PARSE Bioscience Evercode Fixation Kit according to the manufactures protocol. Cells were counted directly before storing at -80°C. Libraries were prepared using the Evercode WT kit (Parse Biosciences) following the manufacturers protocol and sequenced on a Illumina NovaSeq X Plus at a desired depth of 50,000 reads per cell.

Parse Biosciences Trailmaker web server was used for data analysis. The mitochondrial content filter threshold was automatically determined for each sample as 3 median absolute deviations above the median. A third-order spline-type fit was used for the number of genes versus transcripts filter, with the prediction interval threshold automatically determined by Trailmaker. Doublets were identified using the scDblFinder algorithm with automatically determined sample-specific thresholds. Data was integrated with Harmony in Trailmaker and uploaded into BioTuring. CD4 T cells were identified by CD3, TCRalpha, and CD4 expression and an absence of CD8 and CD19.

### *In vitro* stimulation of murine CD4 T cells for qPCR

5×10^5^ CD4 T cells were added to a 96-well plate in serum free RPMI media and allowed to rest for 15min at 37°C. 2X IFNβ (PBL Assay Science Catalog #12401) (200units/mL) was added to a final concentration of 100units/mL IFNβ. Cells were placed in a 37°C incubator for the indicated incubation periods. For qPCR four wells were pooled to create one sample. Cells were placed directly into Qiagen Buffer RLT using dilution concentrations according to the manufacturer’s instructions (Catalog #79216).

### *In vivo* IFN-I stimulation and FACS sorting

Uninfected WT and CD6 KO mice were administered recombinant murine IFNβ (PBL assays Catalog # 12401-1) by intraperitoneal injection of 100units in 100uL of sterile 1X PBS or vehicle control in the evening. Spleens were isolated the following morning, and single-cell suspensions were made. After RBC lysis, cells were washed by centrifugation at 450g for 5min at 4°C and resuspended in FACS buffer (1X PBS with 1% BSA). Cells were then treated with Fc block (clone 2.4G2) for 10min at 4°C, followed by staining for 30min at 4°C with APC anti-CD4 (Clone RM4-5), FITC anti-CD3 (clone 17A2), PerCP-Cy5.5 CD8 (clone 53-6.7), and PE anti-CD44 (clone IM7) monoclonal antibodies. Following staining, cells were washed, resuspended in FACS buffer containing Dapi (BD catalog #564907), and sorted using a BD FACSAria II. Cells were gated as Live, CD3^+^ cells. CD4^+^ CD44^-^cells and CD8^+^ CD44^-^cells were sorted directly into Qiagen Buffer RLT using dilution concentrations according to the manufacture’s instructors (Catalog number 79216).

### qPCR

RNA was isolated using the Qiagen RNeasy MicroKit (Catalog #74004) according to the manufacturer’s instructions. Samples were DNaseI treated (Invitrogen Catalog number: 18068015) according to the manufacturer’s instructions and cDNA was synthesized using MMLV reverse transcriptase (Invitrogen Catalog number: 28025021). Quantitative real-time PCR was performed using SYBR Green Master Mix (Fisher A25742) on a 7500 fast real-time PCR system (Applied Biosystems, Foster City, CA). Transcript levels were calculated relative to the levels of the *Gapdh* housekeeping gene using the following cycle threshold (CT) formula:

[umath1]

### Murine qPCR Primers

m*Ifit1* F: CCTTTACAGCAACCATGGGAGA, R: GCAGCTTCCATGTGAAGTGAC, m*Ifi213* F: GATGGAAGCTTGGGAAGTAGAA, R: GAGAGAACGAGCTTAGTGGATG, m*Rsad2* F: GGTGCCTGAATCTAACCAGAAG, R: CCACGCCAACATCCAGAATA, m*Gapdh* F: CATGGCCTTCCGTGTTCCTA, R: ATGCCTGCTTCACCACCTTCT.

### *In vitro* stimulation of human PBMCs

PBMCs were thawed quickly in a 37°C water bath and washed with serum free RPMI. 5×10^5^ cells were plated into a 24-well plate in serum free RPMI media and allowed to rest for 15min at 37°C. Cells were treated with IgG control (BD Biosciences #550878) or anti-CD6 blocking antibody clone UMCD6 (Millipore MABF2105-100UG) for 15min at 37°C^13^. Human IFNβ was then directly added to a final concentration of 10ug/mL antibody^13^ and 100units/mL IFNβ. Cells were incubated at 37°C for 18-20 hours, fixed with 4% PFA, washed, and resuspended in ice cold 90% Methanol for 15min on ice. Cells were then washed in 1X PBS two times and stained for flow cytometry analysis.

### *In vitro* stimulation of murine splenic cells for flow cytometer

5×10^5^ splenic CD4 T cells were plated into a 96-well plate in serum free RPMI media and allowed to rest for 15min at 37°C. 2X IFNβ (PBL Assay Science Catalog #12401) (200units/mL) was added bringing the final concentration of 100units/mL IFNβ. Cells were placed in a 37°C incubator for 15min and then immediately fixed at 37°C by adding prewarmed 4% PFA (final concentration of 2% PFA). Cells were then centrifuged at 450g for 5min at 4°C. Supernatant was removed and cells were resuspended in ice cold 90% Methanol and incubated on ice for 15min. Cells were then centrifuged at 450g for 5min at 4°C and resuspended in 1X PBS two times.

### Flow Cytometry

Cells were treated with Fc block (Murine cells: clone 2.4G2 human cells: BD #564220) for 15min at 4°C. Cells were stained for 30min at 4°C degrees in FACS buffer using: <u>Murine cells including analysis of CD6 on LL171 cells:</u> PE anti-pSTAT1 clone pY701 (BD catalog #612564), PE anti-IFNAR1 (clone MAR1-5A3), PerCP-Cy5.5 anti-CD3 (clone clone 53-6.7), APC anti-CD4 (Clone RM4-5), BV421 anti-CD44 (clone IM7), BV786 anti-CD45 (clone 30-F11), BUV395 anti-CD6 (clone J90-462). <u>Human cells</u>: BV421 or FITC anti-CD4 (clone RPA-T4), PE anti-IRF7 (MNGPKL), PerCP-Cy5.5 anti-CD3 (UCHT1), PE-Cy7 or BV786 anti-CD8 (SK1), BV421 anti-CD44 (clone SK3), Alexa Flour 647anti-Granzyme B (clone GB11), PE Viperin (MaP.VIP), FITC anti-Ki67 rb545, BV395 anti-T-bet 04-466.

Cells were washed in FACS buffer and resuspended in FACS buffer or fixed in 4% PFA. Data was collected using a 6-laser Beckman CytoFLEX LX. The resulting data were compensated and analyzed with FlowJo software (Tree Star, Inc., Ashland, OR).

### Protein-protein interaction predictions

Modeling interactions between the extracellular domains of human CD6 (amino acids 1-401), IFNaR1 (amino acids 1-436), and IFNaR2 (amino acids 1-244) was first predicted using AlphaFold3 with default settings^35–37^. The top-ranked models were exported as PDB files and analyzed in PyMOL to assess potential interfaces. The highest-ranked CD6–IFNAR1–IFNAR2 model was subjected to all-atom molecular dynamics simulations using GROMACS (version 2023.1)^38–40^. The complex was prepared using the CHARMM27 force field and solvated in a SPC216 water within a cubic simulation box, with counterions added to neutralize the system. Energy minimization was performed using steepest descent algorithm followed by equilibration under constant volume (NVT) conditions for 100 ps and constant pressure (NPT) conditions for 100ps at 300 K and 1 bar. A 100-ns production simulation was subsequently performed using a 2-fs time step with periodic boundary conditions. Temperature and pressure were maintained using the V-rescale thermostat and Parrinello-Rahman barostat, respectively. Long-range electrostatic interactions were calculated using the Particle Mesh Ewald (PME) method with a cutoff distance of 1.0 nm. Covalent bonds involving hydrogen atoms were constrained using LINCs algorithm. Trajectory analyses, including root mean square deviation (RMSD), radius of gyration (Rg), and structural visualization, were performed using standard GROMACS analysis tools to assess conformational stability and persistence of the predicted protein–protein interactions throughout the simulation.

### LL171 Cell Luciferase assay

LL171 cells were cultured in DMEM supplemented with 10% FCS, L-Glutamine, and Penn/Strep and split using trypsin. Experiments were carried out on cells from passage 3-6. 2.5×10^5^ cells were plated into each well of a 96-well plate and incubated at 37°C for around 6hrs to allow the cells to adhere. Cells were washed with serum free media. Recombinant mouse sCD6 was added to the cells for 10min prior to addition of recombinant mouse IFNβ (R&D System #9946-CD-100) for a final concentration of 50ug/mL of sCD6 and 100untis/mL of IFNβ unless otherwise satated. Cells were incubated for 18 hours at 37°C, washed two times with 1X PBS, lysed, and analyzed for luciferase using the Pierce Firefly Luciferase Glow Assay Kit (ThemoFisher #16176) with or without Pierce Firefly Signal Enhancer to increase the signal (ThermoFisher #16180) using a BioTek SYNERGY H1 microplate reader. CD6Δd1 and d1CD6 were custom proteins purchased from GenScript. For these experiments, the buffer the proteins were received in was used to dilute control sCD6 to the same concentration. To control for differences in protein size a final concentration of sCD6, CD6Δd1, and d1CD6 was 100nM. The sequence of CD6 is provided with the sequence for sCD6 in bold and the sequence of d1CD6 versus CD6Δd1 being not underlined versus underlined respectively.

MWLFLGIAGLLTAVLSGLPSPAPSGQHKNGTIPNMTLDLEERLGIRLVNGSSRCSGSV KVLLESWEPVCAAHWNRAATEAVCKALNCGDSGKVTYLMPPTSELPPGATSGNTS SAGNTTWARAPTERCRGANWQFCKVQDQECSSDRRLVWVTCAEN<u>QAVRLVDGSSRCAGRVEMLEHGEWGTVCDDTWDLQDAHVVCKQLKCGWAVKALAGLHFTPGQGPIHRDQVNCSGTEAYLWDCPGRPGDQYCGHKEDAGVVCSEHQSWRLTGGIDSCEGQVEVYFRGVWSTVCDSEWYPSEAKVLCRSLGCGSAVARPRGLPHSLDGRMYYSCKGQEPALSTCSWRFNNSNLCSQSRAARVVCSGSQRHLNLSTSEVPSRVPVTIESSVPVSVKDKDSQG</u>LTLLILCIVLGILLLVSTIFIVILLLRAKGQYALPASVNHQQLSTANQAGINNYHPVPITIAKEAPMLFIQPRVPADSDSSSDSDYEHYDFSSQPPVALTTFYNSQRHRVTEEEA QQNRFQMPPLEEGLEELHVSHIPAADPRPCVADVPSRGSQYHVRNNSDSSTSSEEGYCND PSSKPPPWNSQAFYSEKSPLTEQPPNLELAGSPAVFSGPSADDSSSTSSGEWYQNFQPPPQ HPPAEQFECPGPPGPQTDSIDDDEEDYDDIGAA

### DuoLink Flow-Based Proximity Ligation Assay^44–45^

Protein-protein associations were assessed using the DuoLink® Flow-Based Proximity Ligation Assay (PLA) kit (Sigma DUO94102) according to the manufacturer’s protocol with minor modifications. Primary human PBMCs were fixed with 2% PFA, blocked, and incubated with primary antibodies as indicated in the results section at 1:200 for 60min at 4°. DuoLink PLA probes, ligation, amplification, and detection steps (60min option used) were carried out according to the manufacturer’s instructions. To maintain non-permeabilized conditions and preserve cell surface interactions, all wash steps were performed in the absence of permeabilization and 1× PBS was used in place of the manufacturer’s wash buffer. Data was collected using a 6-laser Beckman CytoFLEX LX and analyzed with FlowJo software (Tree Star, Inc., Ashland, OR).

For murine IFNaR1-IFNaR2 association, splenic CD4 T cells were isolated from UI mice as described above. Isolated CD4 T cells were fixed in 4% PFA for 30 minutes at room temperature. Cells were washed twice in 1X PBS and stained for 30minustes at 4°C using the anti-mouse IFNaR1 clone A5-A3 and IFNaR2 close EPR30040-606 in FACS buffer. Antibody was washed twice in 1X PBS and stained with Duolink probes for 30 minutes at room temperature in 1X PBS as done above. Ligation, amplification, and detection were carried out according to manufacturer’s protocol and outlined as above. For this experiment data was collected on a BD LSRFortessa.

### BiFC

HEK293T cells were co-transfected with 2^nd^ generation lentiviral packaging vectors and lentiviral expression constructs encoding: 1) human CD6 fused to the N-terminal Venus fragment (CD6Venus-N), 2) human STAT1 fused to the C-terminal Venus fragment (STAT1Venus-C), or 3) human IFNAR2 fused to the C-terminal Venus fragment (IFNAR2Venus-C) using Polyethylenimine. For IFNAR2 interaction studies, HEK293T were cells stably transduced with CD6Venus-N and IFNAR2Venus-C constructs and selected using puromycin (1ug/mL) and Hygromycin B (200 ug/mL). Cells were fixed with 4% PFA, and where indicated, immunostained with anti-STAT1 ([Cell Signaling, #9172]) followed by an appropriate fluorescent secondary antibody. Venus fluorescence and STAT1 localization were imaged using a Leica DMi8 Confocal microscope with a 63X objective. For STAT1 interaction studies, HEK293T were cells stably transduced with CD6Venus-N and STAT1Venus-C constructs and selected using puromycin (1ug/mL) and Hygromycin B (200 ug/mL). Cells were fixed with 4% PFA and Venus fluorescence was imaged using a Keyence BZ-X700 with a 40X objective.

Images were analyzed using Image J software (NIH; http://rsbweb.nih.gov/ij) implementing the FIJI plugin set (http://pacific.mpi-cbg.de/wiki/index.php/Fiji).

### Mass Spec analysis

Mass Spec data of an affinity tag pull down from primary murine CD4 T cells transduced to express WT or affinity tagged CD6 was performed by Mori *et al*.^7^ The fold change in the prey protein pulled down in affinity tagged over WT CD6 at time 0 (no TCR re-stimulation) was graphed versus the P value (determined by two-tailed Welch T test).

### Bulk RNAseq

Data files for GSE195543^46^ were downloaded from the NCBI GEO database. Adapter sequences and poor-quality reads were removed followed by use of FastQC to confirm quality control. Paired-reads were aligned and mapped to the human genome GRCh38 using HISAT2. DEG analysis was done using Bioconductor DESeq2.

### Lymphoid Organoids

Splenic lymphoid organoids were generated as described previously^65^. Briefly, 100uLs of splenic single-cell suspensions at a concentration of 6×10^7^ cells/mL was added to transwell inserts (Corning #3460). Inserts were added to 12 well plates with 1mL of prewarmed organoid media supplemented with 1ug/mL of human BAFF (BioLegened 559608)^65^. After resting cells for 30mintures at 37°C, anti-CD6 UMCD6 or IgG control (InvivoGen mabg1-ctrlm) (final concentration of 10ug/mL) was added to the transwell insert followed by 1ug of HA in adjuvant (InvivoGen vac-imq and vac-phos-250)^65^. Every 2-3 days media in the outerwell was supplemented with fresh media containing human BAFF (Biolegend 559608) and UMCD6 or control IgG. On day 7 PI cells were isolated from the transwell and flow cytometry was carried out as described above.

### Statistics

GraphPad Prism software was used to plot data points with SEM for error bars. Statistical significance was determined using GraphPad Prism Software as specified in the figure legends. For scSeq data BioTuring was utilized where Wilcox tests were carried out.

## Resource Availability

## Material availability

This study did not generate new reagents; all reagents will be made available on request after completion of a materials transfer agreement (MTA).

## Data and Code Availability

This paper does not include original code. The bulk RNAseq data was obtained from GSE195543 and COVID-19 scSeq from GSEA 155223. MassSpec data was obtained from the proteomeXchange Consortium via the PRIDE partner repository under the dataset PXD018527. Any additional information required to reanalyze the data reported in this paper is available from the lead contact. Upon manuscript acceptance, our mCoV scSeq RNA sequencing data will be deposited in the Gene Expression Omnibus public repository under the accession number GEO:pending.

## Acknowledgements

scSeq was carried out in combination with the Center for immunotherapy & Precision Immuno-Oncology Immunomonitoring Lab.

This work was supported by the NIH R01 NS086299 (C.B.), NIH R01 EY031463 (F.L.), and NIH R01 HL166544 (Y.W.).

## Author contributions

**A.C-B.** designed the overall study, performed assays, data analysis, data interpretation, and wrote the paper with all co-authors. **A.M.B.** designed the overall study, performed assays, performed scSeq and bulk RNA seq bioinformatics, data analysis, **R. Z.** performed assays, **R.S**. participated in study analysis, supervision, and funding, **F.L.** participated in study design, data interpretation, and funding, **B.T.G.** participated in study design and data interpretation, **Y.W.** designed the overall study, performed assays, data interpretation, supervision, and funding, **C.C.B.** designed the overall study, data interpretation, supervision, and funding.

---

**Supplemental Figure 1:**
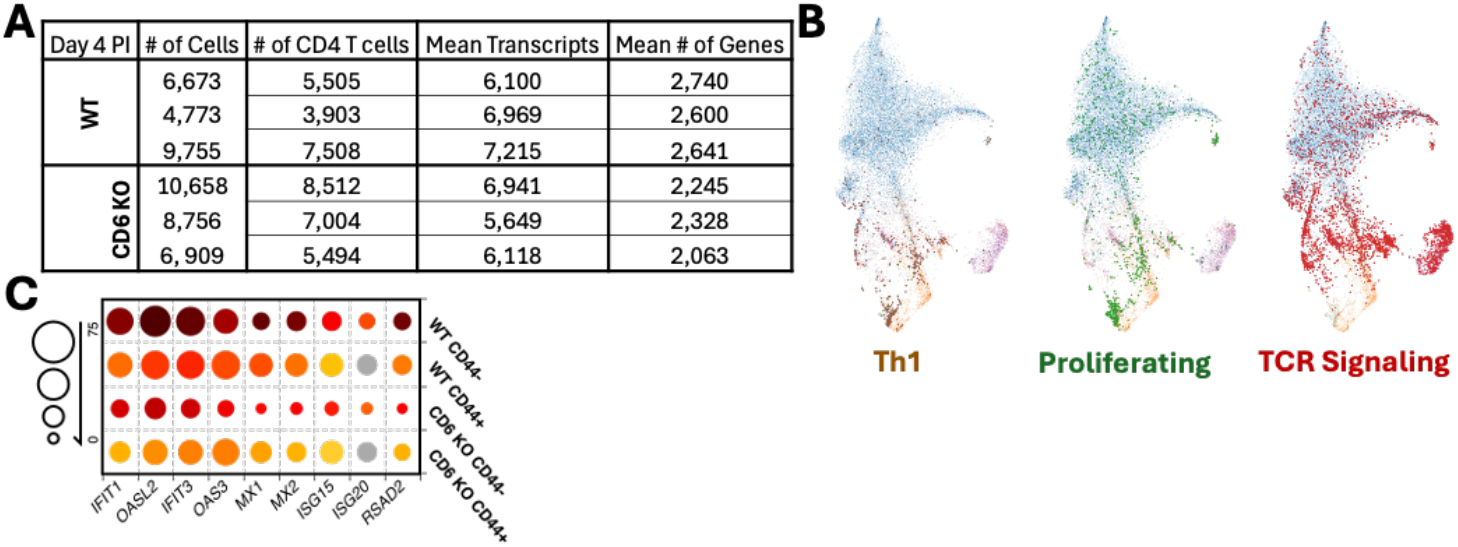
scSeq of CD4 T cells isolated on Day 4 post infection with mCoV encephalomyelitis. **A)** Cell counts and transcript data used for scSeq analysis of CD4 T cells from the dLN on day 4 PI. Each row represents an individual sample obtained from a single mouse. **B)** UMAP highlighting the Th1 (left), proliferating (middle), and TCR signaling (right) clusters. **C)** Heatmap of ISG expression in WT and CD6 KO CD4 T cells at day 4 PI separated by *Cd44* expression.

**Supplemental Figure 2:**
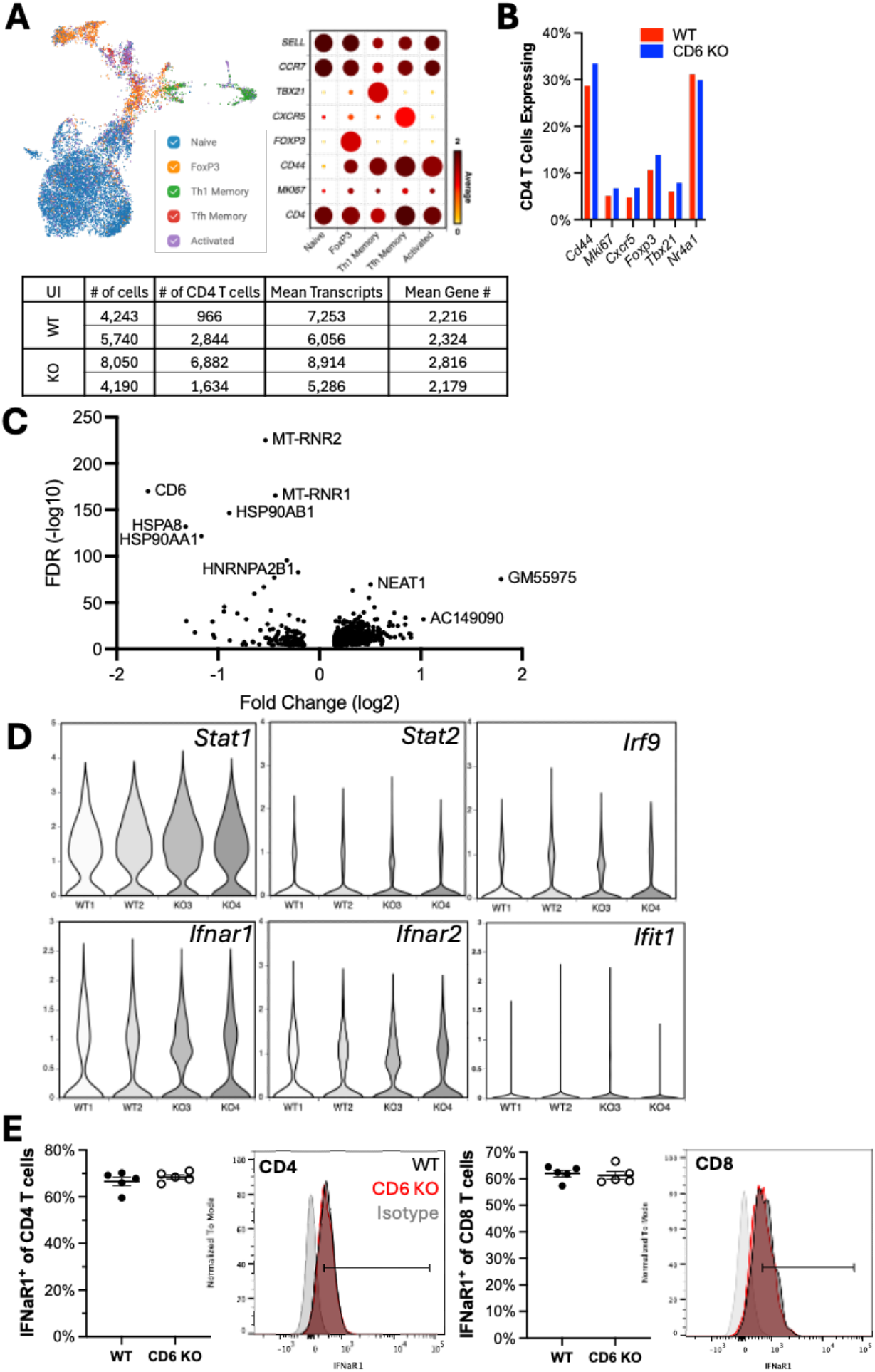
CD6 deficient T cells exhibit largely preserved homeostatic features. scSeq was performed on CD4 T cells negatively enriched from uninfected (UI) WT and CD6 KO mice. **A)** UMAP showing CD4 T cell clusters from UI mice (left) and identified by established markers (middle). Cell counts and transcripts of each WT and CD6 KO sample was provided with each row represents an individual sample obtained by pooling two mice. **B)** Distribution of clusters between UI WT and CD6 KO CD4 T cells was graphed as the percent of cells positive for identifying effector markers (right) and **C)** DEGs were plotted on a volcano plot. **D)** Transcription of IFN-I signaling components were plotted and **E)** comparable cell surface IFNaR1 protein levels in T cells from UI spleens were confirmed by flow cytometry.

